# Direct and indirect RANK ligand and CD40 ligand signaling regulate the maintenance of thymic epithelial cell frequency and properties in the adult thymus

**DOI:** 10.1101/2024.10.10.617559

**Authors:** Mio Hayama, Hiroto Ishii, Maki Miyauchi, Masaki Yoshida, Naho Hagiwara, Wataru Muramtatu, Kano Namiki, Rin Endo, Takahisa Miyao, Nobuko Akiyama, Taishin Akiyama

## Abstract

Medullary thymic epithelial cells (mTECs) play a crucial role in suppressing the onset of autoimmunity by eliminating autoreactive T cells and promoting the development of regulatory T cells in the thymus. Although mTECs undergo turnover in adults, the molecular mechanisms behind this process remain unclear. This study describes the direct and indirect roles of receptor activator of NF-κB ligand (RANKL) and CD40 ligand (CD40L) signaling in TECs in the adult thymus. Flow cytometric and single-cell RNA-seq (scRNA-seq) analyses suggest that the depletion of both RANKL and CD40L signaling inhibits mTEC differentiation from CCL21^+^ mTEC progenitors to transit-amplifying TECs in the adult thymus. Unexpectedly, this depletion also indirectly affects the gene expression of TEC progenitors and cortical TECs. Notably, AP-1 gene expression, which allows further subdivision of TEC progenitors, is upregulated following the depletion of RANKL and CD40L signaling. Overall, our data propose that RANKL and CD40L signaling cooperatively maintain mature mTEC frequency in the adult thymus and sustain the characteristics of TEC progenitors through an indirect mechanism.

## Introduction

Thymic epithelial cells (TECs) are required for the differentiation of self-tolerant T cells and regulatory T cells in the thymus. TECs are separated into cortical TECs (cTECs) and medullary TECs (mTECs) depending on their localization in the thymus ^1^. In addition, each TEC subset has distinct properties and functions in T cell selection and differentiation. cTECs are critical for early T cell development and positive selection of thymocytes expressing both surface makers CD4 and CD8. In contrast, mTECs ectopically express tissue-restricted self-antigens (TSAs) to filter out a wide range of self-antigen reactive T cells by apoptosis or to convert them into regulatory T cells. The TSA expression in mTECs is regulated by transcriptional regulator AIRE, which is highly expressed in mTECs expressing high levels of MHC class II and co-stimulatory molecules.

During embryonic development, both mTEC and cTEC differentiate from common bipotent progenitor cells ^2,3^. For mTEC development, claudine 3 and 4-positive TECs ^4^, *Krt19*-positive mTECs ^5^, *Tnfrsf11a*-positive TECs ^6^, *Ccl21a*-positive TECs ^7^, and *Pdpn*-expressing TECs ^8^ were reported as mTEC progenitors giving rise to mTECs expressing AIRE and TSAs. In the adult thymus, Aire^+^ mTECs undergo a turnover of approximately 2 weeks ^9^, indicating the presence of mTEC progenitor maintaining the cellularity of mature mTECs. Some studies propose the progenitor of TECs in the adult thymus ^10,11^. However, the phenotypes of the proposed progenitors seem to be inconsistent, implying that multiple fractions of TECs may have the potential as TEC progenitors.

Single-cell RNA-sequencing (scRNA-seq) analysis is a powerful tool for distinguishing cell types with high resolution. Recent studies utilizing scRNA-seq on TECs have highlighted their significant heterogeneity. Beyond identifying AIRE^+^ mTECs and CCL21^+^ mTECs, data analysis has revealed the presence of transit-amplifying TECs (TA-TECs) ^12–15^, which are proliferative progenitors for AIRE^+^ mTECs, as well as post-AIRE mTECs, including tuft-like mTECs and mimetic TECs ^16,17^. Additionally, a recent study suggested the existence of TEC progenitors expressing a wide variety of keratin molecules in the human thymus ^18^. Moreover, a combination of scRNA-seq analysis and barcode cell labeling has proposed the presence of the early and late types of TEC progenitors in postnatal mice ^19^ although these TEC progenitors have not been isolated and fully characterized yet.

Mechanistically, several studies have revealed the roles of TNF family cytokine signaling in mTEC differentiation. Receptor activator of NF-κB (RANK) ligand and CD40 ligand play partially redundant roles in mTEC differentiation during early thymic development by activating signal transducer TRAF6- and NF-κB inducing kinase-dependent activation of transcription factor NF-κB ^20–22^. Additionally, lymphotoxin signaling is involved in early mTEC differentiation by inducing the expression of RANK on embryonic mTEC progenitors ^6,23^, postnatal development of CCL21^+^ mTECs ^24^, and differentiation of post-Aire mTECs ^25^. The administration of a RANKL neutralizing antibody (RANKL-Ab) results in a reduction of mature mTECs ^13,26^, suggesting that RANKL signaling is involved in the homeostatic maintenance of AIRE^+^ mTEC frequency in the adult thymus.

Activator Protein 1 (AP-1) is a family of dimeric transcription factors including JUN, FOS, ATF, and MAF family members. AP-1 is activated by various stimuli, including cytokines and growth factors, through mitogen-activated protein kinase (MAPK) cascades, and thereby regulates numerous cellular and physiological functions ^27^. In a study of TEC development, FOS expression driven by the H2-Kb promoter was shown to cause thymic hyperplasia by expanding TECs ^28^. Additionally, RANKL signaling can activate the MAPK cascade via TRAF6 ^29^, a signal transducer critical for mTEC differentiation ^30^, implying a possible role for AP-1 in this process.

In this study, we describe how RANKL and CD40L signaling redundantly support the differentiation of CCL21+ mTECs into TA-TECs, thereby maintaining the frequencies of Aire+ mTECs and Post-Aire mTECs in the postnatal thymus. Unexpectedly, depletion of both RANKL and CD40L signaling also has indirect effects on the gene expression profiles of TEC progenitors and cortical TECs.

Additionally, after the depletion of RANKL and CD40L signaling, the expression levels of AP-1 genes, which facilitate further subdivision of TEC progenitors, are up-regulated. Overall, our data suggest that these TNF family cytokine signals directly and indirectly regulate TEC frequency and properties.

## Results

### RANK and CD40 signaling redundantly maintain mature mTEC cellularity

Aligned with the reported role of RANKL signaling in maintaining mature mTECs in adult mice ^26^, flow cytometric analysis confirmed that blocking RANKL-RANK signaling with an anti-RANKL antibody (RANKL-Ab) significantly reduces the number of mTECs expressing high MHC class II (MHCII^hi^UEA-1^+^Ly51^−^TECs; mTEC^hi^) two weeks after the administration in mice (WT-RANKL Ab) compared to control IgG administration (WT-Control) (Figure 1A and B). However, approximately 10% of the mTEC^hi^ population persisted in the thymus of WT-RANKL Ab (Figure 1A and B). We speculated that CD40 signaling might compensate for the absence of RANKL signaling in maintaining adult mTECs, similar to its role during mTEC development in embryonic and neonatal stages ^20^. To test this hypothesis, we administered RANKL-Ab to *Cd40*-deficient mice (*Cd40*^−/−^-RANKL Ab). Indeed, neutralizing RANKL signaling in *Cd40*^−/−^ mice resulted in a reduction of mTEC^hi^ cell numbers to just a few percent of those in WT-RANKL Ab and *Cd40*- deficient mice receiving control IgG (*Cd40*^−/−^-Control) (Figure 1A and B). In contrast to mTEC fractions, cell numbers of Ly51^+^UEA^−^ TECs (cTECs) and Ly51^−^UEA^−^TECs were unaffected by the RANKL-Ab administration and the *Cd40*-deficiency. These results suggest that RANKL and CD40L signaling contribute to maintaining the frequency of mature mTECs in the adult thymus in a partially redundant manner, but not the frequency of other TECs.

**Figure 1.**
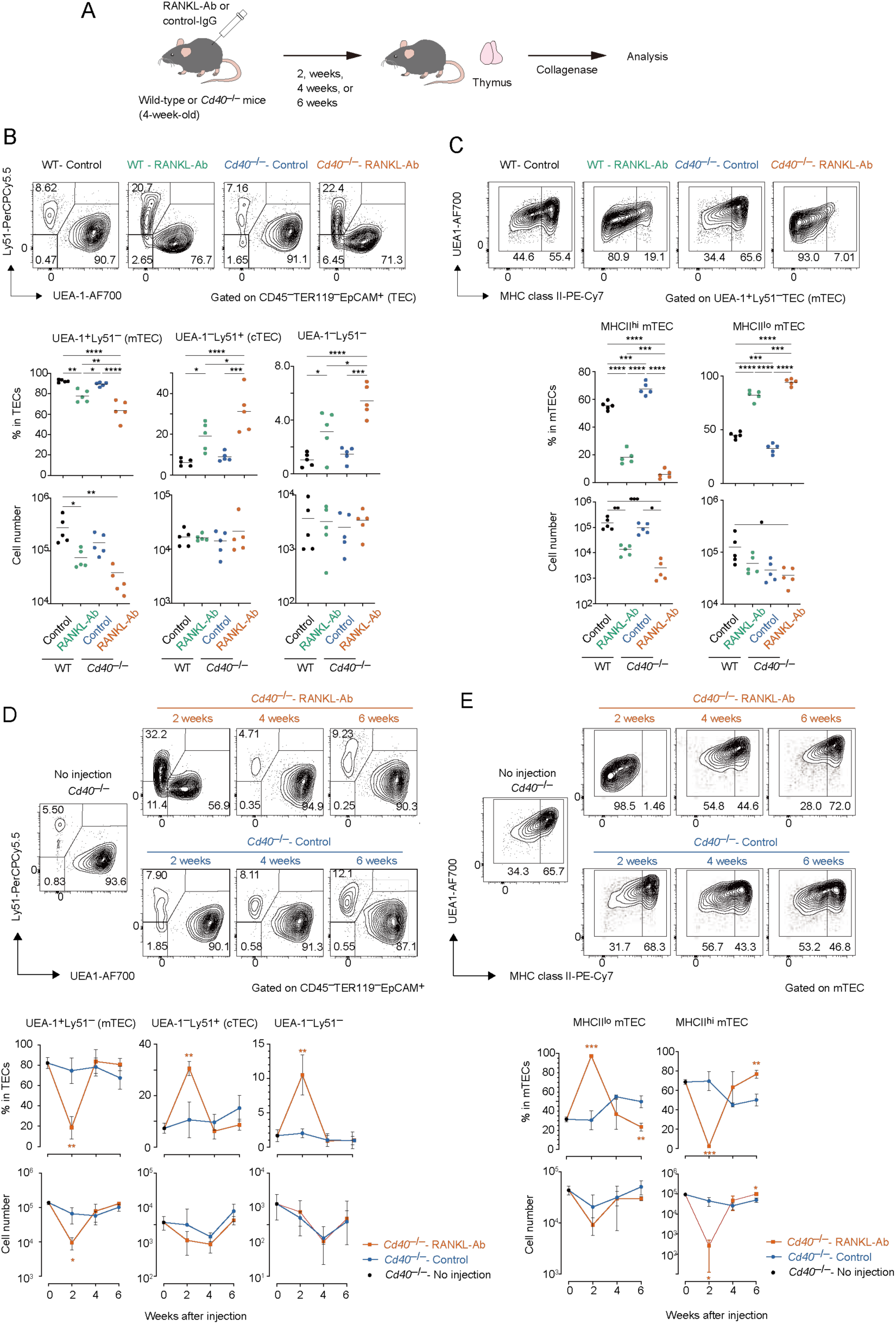
Flow cytometric analysis of TECs from wild-type and *Cd40*-deficient mice treated with neutralizing RANKL antibody. (A) Experimental scheme for depleting RANKL signaling by the administration of RANKL antibody in mice. (B) Flow cytometric analysis of UEA-1 ligand and Ly51 expressions in TECs (CD45^−^Ter-119^−^ EpCAM^+^) from wild-type (WT) treated with control IgG (WT-Control), WT treated with neutralizing RANKL antibody (WT-RANKL-Ab), *Cd40*-deficient (*Cd40*^−/−^) mice treated with control-IgG (*Cd40*^−/−^-Control), and *Cd40*^−/−^ mice treated with RANKL-Ab (*Cd40*^−/−^-RANKL-Ab) at 6-week-old age (n = 5 each). The percentages and numbers of UEA-1^+^Ly51^−^ (mTEC), UEA-1^−^Ly51^+^ (cTEC), and UEA-1^−^Ly51^−^ in TECs are summarized in graphs. RANKL-Ab or control IgG was subcutaneously injected in mice at 4-week-old age. Bars indicate the mean value. Data were statistically analyzed using one-way ANOVA followed with multiple comparisons by Tukey’s test. Significant differences are indicated by * p < 0.05, ** p < 0.01, *** p < 0.001, **** p < 0.0001. (C) Flow cytometric analysis of MHC class II (MHCII) and UEA-1 ligand expressions in mTECs (UEA-1^+^Ly51^−^ TECs) from WT-Control, WT-RANKL-Ab, *Cd40*^−/−^-Control, *Cd40*^−/−^-RANKL-Ab at 6-week-old age (n = 5). The percentages and numbers of MHCII^hi^UEA-1^+^ cells and MHCII^lo^UEA-1^+^ mTEC in mTECs are summarized in graphs. Bars indicate the mean value. Data were statistically analyzed using one-way ANOVA followed with multiple comparisons by Tukey’s test. Significant differences are indicated by* p < 0.05, ** p < 0.01, *** p < 0.001, **** p < 0.0001. (D) Flow cytometric analysis of UEA-1 ligand and Ly51 expressions in TECs from *Cd40^−/−^* mice 2, 4, and 6 weeks after the treatment with RANKL-Ab (*Cd40*^−/−^-RANKL-Ab) or control IgG (*Cd40*^−/−^- Control), and no treatment (no injection). N = 3 each. The percentages and numbers of UEA- 1^+^Ly51^−^ (mTEC), UEA-1^−^Ly51^+^ (cTEC), and UEA-1^−^Ly51^−^ in TECs are summarized in graphs. Data are expressed as the mean ± SD. Data were statistically analyzed using unpaired t-test. Significant differences are indicated by * p < 0.05, ** p < 0.01. (E) Flow cytometric analysis of MHC class II (MHCII) and UEA-1 ligand expressions in mTECs from *Cd40^−/−^* mice 2, 4, and 6 weeks after the treatment with RANKL-Ab (*Cd40*^−/−^-RANKL-Ab) or control IgG (*Cd40*^−/−^-Control), and no treatment (no injection). N = 3 each. MHCII^hi^UEA-1^+^ cells and MHCII^lo^UEA-1^+^ mTEC in mTECs are summarized in graphs. Bars indicate the mean value. Data are expressed as the mean ± SD. Data were statistically analyzed using unpaired t-test. Significant differences are indicated by * p < 0.05, ** p < 0.01, *** p < 0.001.

Although mTEC^hi^ was severely reduced in the thymus of *Cd40*^−/−^-RANKL-Ab, mTECs expressing low levels of MHC class II (mTEC^lo^) were less affected. The number of mTEC^lo^ cells was reduced by approximately half, with a substantial number remaining in the thymus of *Cd40*^−/−^-RANKL-Ab. Given that the mTEC^lo^ fraction includes immature mTECs in addition to post-Aire mTECs ^31^, it is likely that precursors for mature mTECs persist in the mTEC^lo^ fraction in these mice. Indeed, mature mTECs were restored in *Cd40*^−/−^-RANKL-Ab 4 weeks after RANKL administration, likely due to the homeostatic clearance of the injected antibody (A 1C and D). Moreover, 6 weeks after administration, the ratio of mTEC^lo^ to mTEC^hi^ shifted; the proportion of mTEC^lo^ decreased while the proportion of mTEC^hi^ increased in total mTECs compared to age-matched controls. This observation supports the idea that the mTEC^lo^ pool serves as a precursor for mTEC^hi^ during the rapid recovery, leading to a reduction in the relative proportion of mTEC^lo^. Overall, these data suggest that immature mTECs remain in the mTEC^lo^ fraction in *Cd40*^−/−^-RANKL-Ab 2 weeks after antibody administration and differentiate into mTEC^hi^ following the clearance of RANKL-Ab

### RANKL and CD40L signaling up-regulate cell-cycle related genes and down-regulates Ccl21a expression in mTEC^lo^ fraction

Given that the mTEC^lo^ fraction remaining after the depletion of RANKL and CD40L signaling might represent the phenotype of mTEC progenitors prior to receiving these cytokine signals, we aimed to investigate the gene expression profile of a specific subfraction of mTEC^lo^ cells in *Cd40*^−/−^-RANKL-Ab. To minimize contamination from post-Aire mTECs, we selectively sorted cells within the mTEC^lo^ fraction that were negative for Ly6d (a marker for post-Aire mTECs) and L1CAM (a marker for tuft-like TECs) ^32^ (Supplementary Figure 1A). These sorted cells were then subjected to RNA sequencing (RNA-seq) analysis to elucidate their gene expression profiles. Principal component analysis (PCA) of the RNA-seq data showed that the gene expression profiles of the mTEC^lo^ subfraction differed significantly among wild-type (WT), WT-RANKL-Ab, *Cd40*^−/−^, and *Cd40*^−/−^- RANKL-Ab mice (Supplementary Figure 1B). Differentially expressed genes (DEGs) were identified using a threshold of a 2-fold change with an FDR P-value < 0.05 (Figure 2A and Supplementary Table 1). The administration of RANKL-Ab to wild-type (WT) mice led to the up-regulation of 111 genes and down-regulation of 274 genes. The deletion of CD40 resulted in the up-regulation of 255 genes and down-regulation of 277 genes. Notably, administering RANKL-Ab to *Cd40*^−/−^ mice induced the up-regulation of 313 genes and down-regulation of 819 genes compared to WT-RANKL-Ab mice. Venn diagram analysis of the DEG sets revealed a significant reduction of 492 genes specifically in *Cd40*^−/−^-RANKL-Ab mice (Figure 2B). These results highlight the redundant and additive effects of RANKL and CD40L signaling in regulating gene expression in immature mTECs.

**Figure 2.**
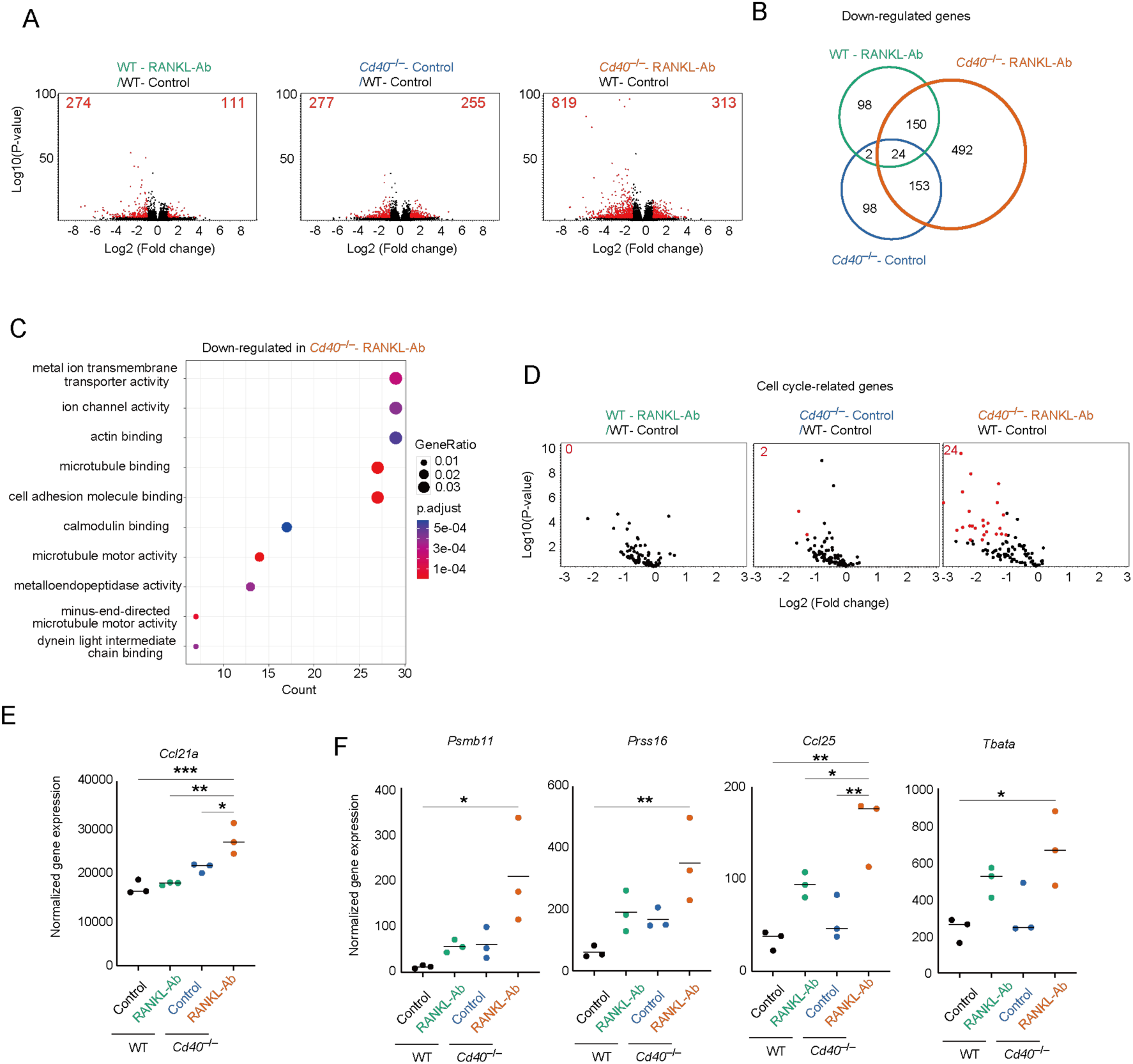
RNA-seq analysis of mTEC^lo^ fraction from wild-type and *Cd40*-deficient mice receiving neutralizing RANKL antibody. (A) Volcano plots of differentially expressed genes from bulk RNA-seq data of wild-type (WT) treated with control IgG (WT-Control), WT treated with neutralizing RANKL antibody (WT-RANKL-Ab), *Cd40*-deficient (*Cd40*^−/−^) mice treated with control-IgG (*Cd40*^−/−^-Control), and *Cd40*^−/−^ mice treated with RANKL-Ab (*Cd40*^−/−^-RANKL-Ab) at 6-week-old age. Red dots in volcano plots indicate genes for which expression differed significantly between the two samples (FDR P-value < 0.05, Fold change > 2). Numbers of differentially expressed genes are shown in the panels. The log2 fold change is plotted on the x-axis, and the log10 P-value is plotted on the y-axis. P-values were determined by Baggerley’s test ^38^ (B) The Venn diagram illustrates the overlap of down-regulated genes among three samples compared to WT mice treated with control-IgG. (C) Gene ontology enrichment analysis of the down-regulated genes in *Cd40^−/−^* mice treated with RANKL-Ab compared to WT mice treated with control-IgG. (D) Volcano plots of differential expression of cell cycle-related gene sets. Red dots in volcano plots indicate genes for which expression differed significantly between the two samples (FDR P-value < 0.05, Fold change > 2). For cell cycle-related gene sets, mouse orthologues of the previously reported human cell cycle-related gene sets ^39^ were used. Numbers of differentially expressed genes are shown in the panels. The log2 fold change is plotted on the x-axis, and the log10 P-value is plotted on the y-axis. P-values were determined by Baggerley’s test ^38^ (E) Dot plot showing normalized gene expression value of *Ccl21a*. The horizontal lines show the mean. Data were statistically analyzed using one-way ANOVA followed with multiple comparisons by Tukey’s test. Significant differences are indicated by * p < 0.05, ** p < 0.01, *** p < 0.001. (F) Dot plots showing normalized gene expression value of some cTEC-associated genes. Data were statistically analyzed using one-way ANOVA followed with multiple comparisons by Tukey’s test. The horizontal lines show the mean. Significant differences are indicated by * p < 0.05, ** p < 0.01.

Gene Ontology (GO) analysis of down-regulated gene sets in the mTEC^lo^ subfraction from *Cd40*^−/−^- RANKL-Ab, compared to WT-control mice, revealed significant enrichment in GO terms associated with ion transport, cytoskeletal organization, and microtubule motor activity (Figure 2C). This suggests that RANKL and CD40L signaling promote the expression of these gene sets in the mTEC^lo^ subfraction. Alternatively, there may be a reduction in the frequency of post-Aire mimetic mTECs that express these gene sets but not L1CAM and Ly6d, potentially influenced by RANKL and CD40L signaling.

In addition to the GO analysis, we found that cell cycle-related gene sets were down-regulated following the disruption of RANKL and CD40L signaling (Figure 2D). Furthermore, *Ccl21a* expression was up-regulated in the mTEC^lo^ subfraction of *Cd40*^−/−^-RANKL-Ab (Figure 2E). These findings suggest that RANKL and CD40L signaling may initiate the differentiation of CCL21^+^ mTECs into transit-amplifying TECs, which serve as precursor cells for Aire^+^ mTECs ^15^. Alternatively, the increased expression of *Ccl21a* in the mTEC^lo^ subfraction might be due to a higher proportion of *Ccl21a*-expressing cells, resulting from a reduction in the frequency of post-Aire mimetic mTECs in this subfraction.

Additionally, genes typically associated with cTECs, including *Psmb11*, *Prss16*, *Tbata*, and *Ccl25*, were up-regulated in the mTEC^lo^ subfraction of *Cd40^−/−^*-RANKL-Ab mice (Figure 2F). This observation indicates that RANKL and CD40L signaling help suppress the aberrant expression of certain cTEC genes in mTECs.

### Single-cell RNA-seq analysis suggested that RANKL and CD40L signaling indirectly regulate gene expressions in cTECs and TEC progenitors in the thymus

Given the high heterogeneity of TECs, the mTEC^lo^ subfraction identified by flow cytometric analysis can encompass multiple TEC subsets including post-Aire mTECs as well as immature mTECs. To gain a more comprehensive understanding of the changes in frequency and gene expression profiles of TECs following the depletion of these cytokine signaling, we conducted single-cell RNA sequencing (scRNA-seq) analysis. Droplet-based scRNA-seq was performed on the TEC fraction (EpCAM^+^CD45^−^TER119^−^) isolated from WT-control, WT-RANKL-Ab, *Cd40*^−/−^- control, and *Cd40*^−/−^-RANKL-Ab mice. After the integration of these scRNA-seq data, TEC clusters were defined based on the expression of marker genes (Figure 3A and Supplementary Figure 2). In addition to the relatively well-characterized TEC subsets, we determined a cluster (Cluster 9 in Figure 3A) that appears to correspond to the early TEC progenitor population previously described ^19^, characterized by expression of *Psmb11*, *Prss16* and *Pdpn* (Figure 3B and Supplementary Figure 2).

**Figure 3.**
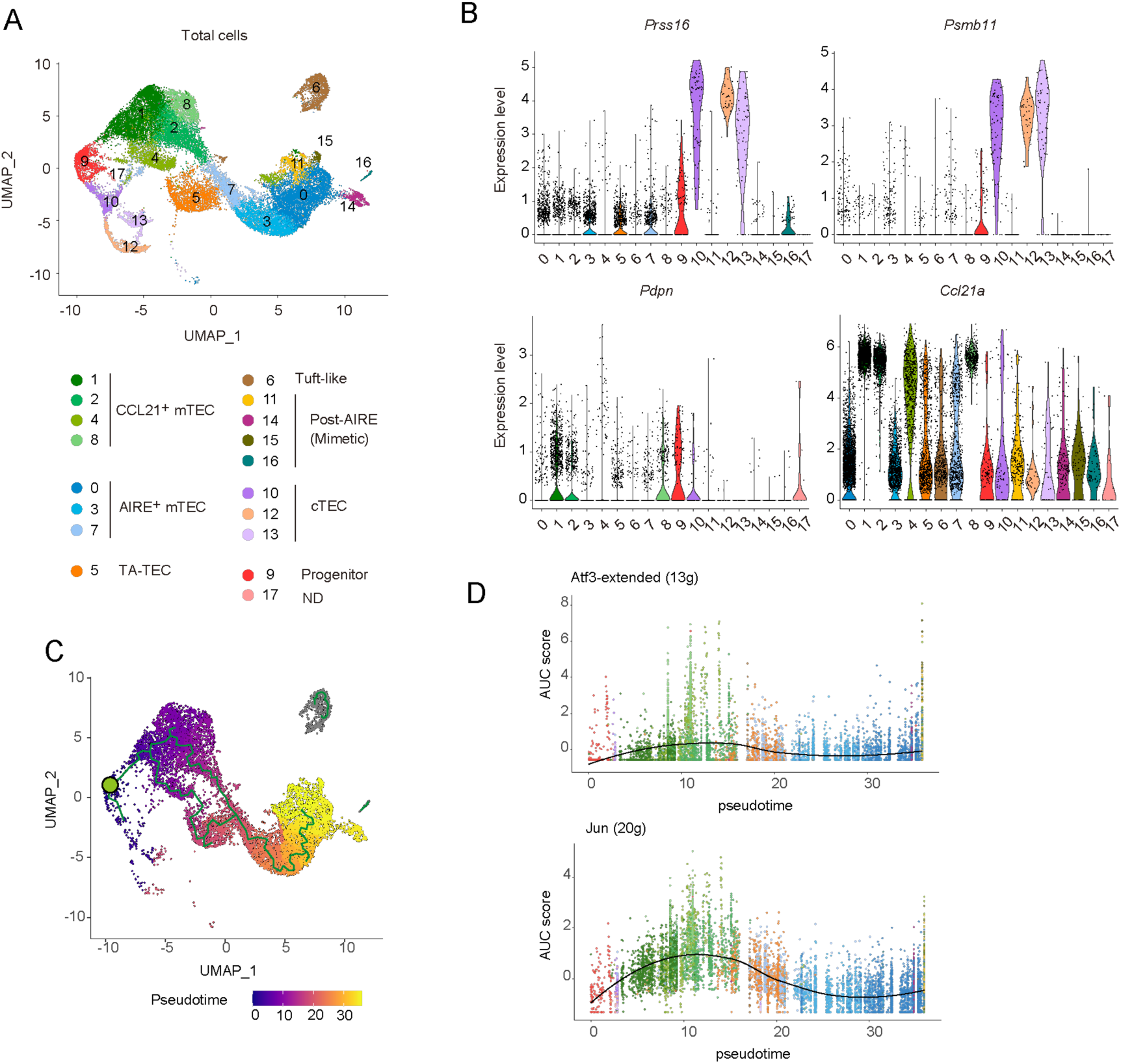
Single RNA-seq analysis of TECs from wild-type and *Cd40*-deficient mice receiving neutralizing RANKL antibody. (A) Uniform Manifold Approximation and Projection (UMAP) plot of droplet-based scRNA-seq data of TECs (CD45^−^Ter-119^−^EpCAM^+^). The scRNA-seq data from wild-type (WT) mice treated with control IgG, WT mice treated with a neutralizing RANKL antibody (RANKL-Ab), *Cd40*- deficient (*Cd40*^−/−^) mice treated with control IgG, and *Cd40*^−/−^ mice treated with RANKL-Ab were integrated using the Seurat package. In the plot, cell clusters are distinguished by colors and numbers and are assigned based on marker gene expression (Supplementary Figure 2) (B) Violin plots showing the expression levels of *Prss16, Psmb11*, *Pdpn*, and *Ccl21a* in each cluster. The expression levels of these genes in WT treated with control IgG were exhibited. Each dot represents expression levels in individual cells. (C) Monocle trajectory and pseudotime analyses of TEC scRNA-seq data from WT treated with control IgG. The green circle indicates the root node when cluster 9 is considered as TEC progenitors. (D) Plot of the area under the recovery curve (AUC), which reflects the enrichment of each regulon, versus pseudotime predicted from Monocle. The regulon of ATF3 and JUN were determined by the SCENIC program.

Monocle trajectory analysis ^33^ using WT-Control TEC clusters indicated that cluster 9 is situated between the CCL21^+^ mTEC and cTEC clusters (Figure 3C), suggesting that this cluster likely represents a progenitor TEC population. When the root node was set in cluster 9 (indicated by the green circle in Figure 3C), a combined analysis using Monocle and SCENIC tools ^34^ demonstrated an increased activity of ATF3- and JUN-associated regulons—groups of genes regulated by shared transcription factors—during the differentiation of progenitor clusters into CCL21^+^ mTECs (Figure 3D and Supplementary Table 2). These findings suggest that the activities of these AP-1 transcription factors may play a role in driving the differentiation of progenitor cells into the mTEC lineage.

Comparison of the scRNA-seq data clusters among WT-control, WT-RANKL-Ab, *Cd40*^−/−^-control, and *Cd40*^−/−^-RANKL-Ab mice revealed a marked reduction in the frequencies of Aire^+^ mTECs, TA-TECs, post-Aire mTECs, and tuft-like mTECs in *Cd40*^−/−^-RANKL-Ab mice (Figure 4A and B). In contrast, RANKL-Ab administration in wild-type mice resulted in a milder reduction of these mTEC subsets (Figure 4A and B). These findings are consistent with those from flow cytometric analysis, further supporting the functional redundancy of RANKL and CD40L signaling in mTEC maintenance in the adult thymus. Additionally, the reduction of TA-TECs in *Cd40^−/−^*-RANKL-Ab (Figure 4A and B) correlates with the down-regulation of cell cycle-related genes in the mTEC^lo^ fraction, identified in the bulk RNA-seq analysis (Figure 2D). This correlation arises because the TA-TEC cluster in scRNA-seq includes not only *Aire*^+^ subcluster but also the *Ccl21a*^+^ subset ^15^, which should be part of the mTEC^lo^ population in flow cytometric analysis. In contrast, the CCL21^+^ mTEC, cTEC, and TEC progenitor clusters appeared to remain in the thymus of *Cd40*^−/−^-RANKL-Ab. Overall, scRNA-seq analysis suggested that RANKL and CD40L signaling redundantly maintain the frequency of Aire^+^ mTECs and post-Aire mimetic mTECs by promoting the differentiation of CCL21^+^ mTEC into TA-TECs in the adult thymus.

**Figure 4.**
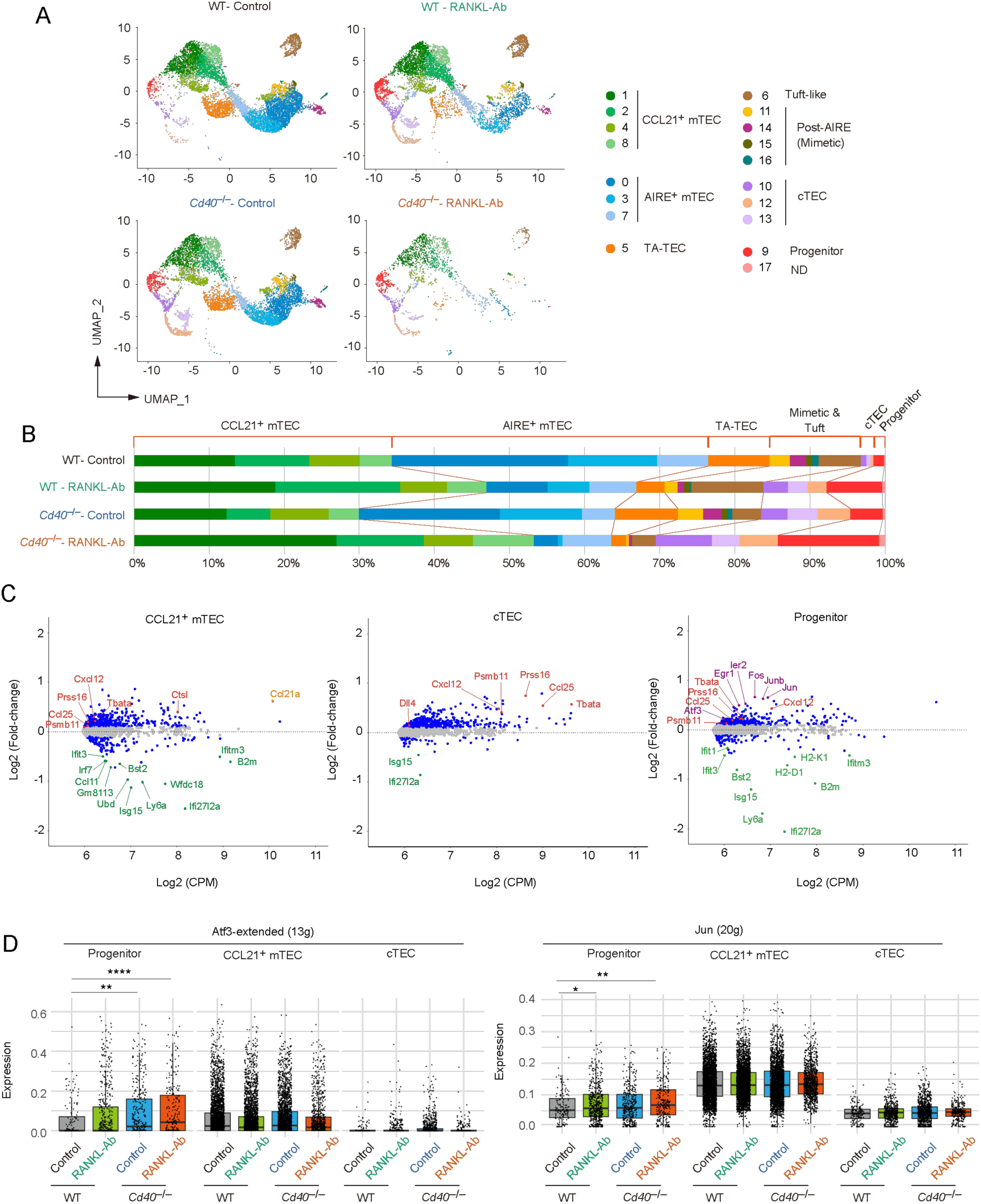
Differentially expressed gene and trajectory analyses of single-cell RNA-seq data from wild-type and *Cd40*-deficient mice receiving neutralizing RANKL antibody. (A) UMAP plot of droplet-based scRNA-seq data of TECs (CD45^−^Ter-119^−^EpCAM^+^) from wild-type (WT) mice treated with control IgG (WT-Control), WT mice treated with a neutralizing RANKL antibody (WT-RANKL-Ab), *Cd40*-deficient (*Cd40*^−/−^) mice treated with control IgG (*Cd40*^−/−^- Control), and *Cd40*^−/−^ mice treated with RANKL-Ab (*Cd40*^−/−^- RANKL-Ab). The integrated UMAP plot in Figure 3 was separated into each data set. (B) Percentages of cell subsets in total TECs were compared among the scRNA-seq data from WT-Control, WT-RANKL-Ab, *Cd40*^−/−^- Control, and *Cd40*^−/−^- RANKL-Ab. (C) MA plots show differentially expressed genes between WT-control and *Cd40*^−/−^- RANKL-Ab from scRNA-seq data for each cell cluster subset. The log2 average expression level (CPM) is plotted on the x-axis, and the log2 fold change is plotted on the y-axis. Genes with log2 fold change greater than 0.1 or less than –0.1 are represented by blue dots. Red dots indicate cTEC-associated genes, violet dots indicate AP-1 transcription factor genes, and green dots indicate interferon-stimulated genes. The orange dot represents Ccl21a. (D) Total expression of ATF3 and JUN regulon genes in the TEC progenitors, CCL21^+^ mTECs, and cTECs subclusters of WT-Control, WT-RANKL-Ab, *Cd40*^−/−^- Control, and *Cd40*^−/−^- RANKL-Ab. ** p < 0.05, ** p < 0.01, and **** p < 0.0001.

In line with findings from bulk RNA-seq analysis, differential gene expression analysis of scRNA-seq data revealed that the depletion of RANKL and CD40L signaling leads to the up-regulation of cTEC-associated genes and *Ccl21a* in CCL21^+^ mTEC clusters (Figure 4C). Additionally, several interferon-stimulated genes (ISGs) were notably down-regulated in these clusters (Figure 4C). Interestingly, these changes in gene expression were observed not only in CCL21^+^ mTECs, which are the primary recipients of RANKL and CD40L signaling, but also in cTECs and TEC progenitors (Figure 4C). Given that the RANK expression of these cell types is virtually absent (Supplementary Figure 3), this unexpected finding suggests an indirect regulatory mechanism of gene expression driven by RANKL and CD40L signaling. Furthermore, the loss of RANKL and CD40L signaling resulted in the up-regulation of some AP-1 transcription factor genes within the progenitor cell subset (Figure 4C). Consistently, the SCENIC analysis suggested an increase in the activity of ATF3- and JUN-inducing regulons in the progenitor TECs (Figure 4D). Collectively, these results imply that under normal conditions, RANKL and CD40L signaling may act to indirectly suppress gene regulatory networks governed by AP-1 transcription factors in progenitor cells, highlighting a complex interplay of direct and indirect signaling pathways in maintaining TEC homeostasis.

### TEC progenitors are classified into subpopulations with unique gene expression profiles

Given that our data suggest TEC progenitors are indirectly influenced by RANKL and CD40L signaling, we focused our analysis on these cells. Subclustering of the progenitor cluster from the scRNA-seq data revealed four distinct subclusters (Figure 5A), all exhibiting similar levels of *Pdpn* expression (Figure 5B). Cluster S1 showed high expression of cTEC-associated genes, such as *Prss16* and *Psmb11*(Figure 5B), suggesting a bias toward the cTEC lineage. In contrast, cluster S2 displayed high levels of *Ccl21a* expression in a part of the cells, indicating a bias toward the mTEC lineage. Clusters S0 and S3 exhibited lower expression levels of both cTEC-associated genes and *Ccl21a* (Figure 5B). Notably, cluster S3 was characterized by the elevated expression of AP-1 transcription factor family genes (Figure 5B and Supplementary Figure 4). These findings underscore the heterogeneous composition of TEC progenitors, categorized by their expression levels of cTEC-associated genes and AP-1 family genes.

**Figure 5.**
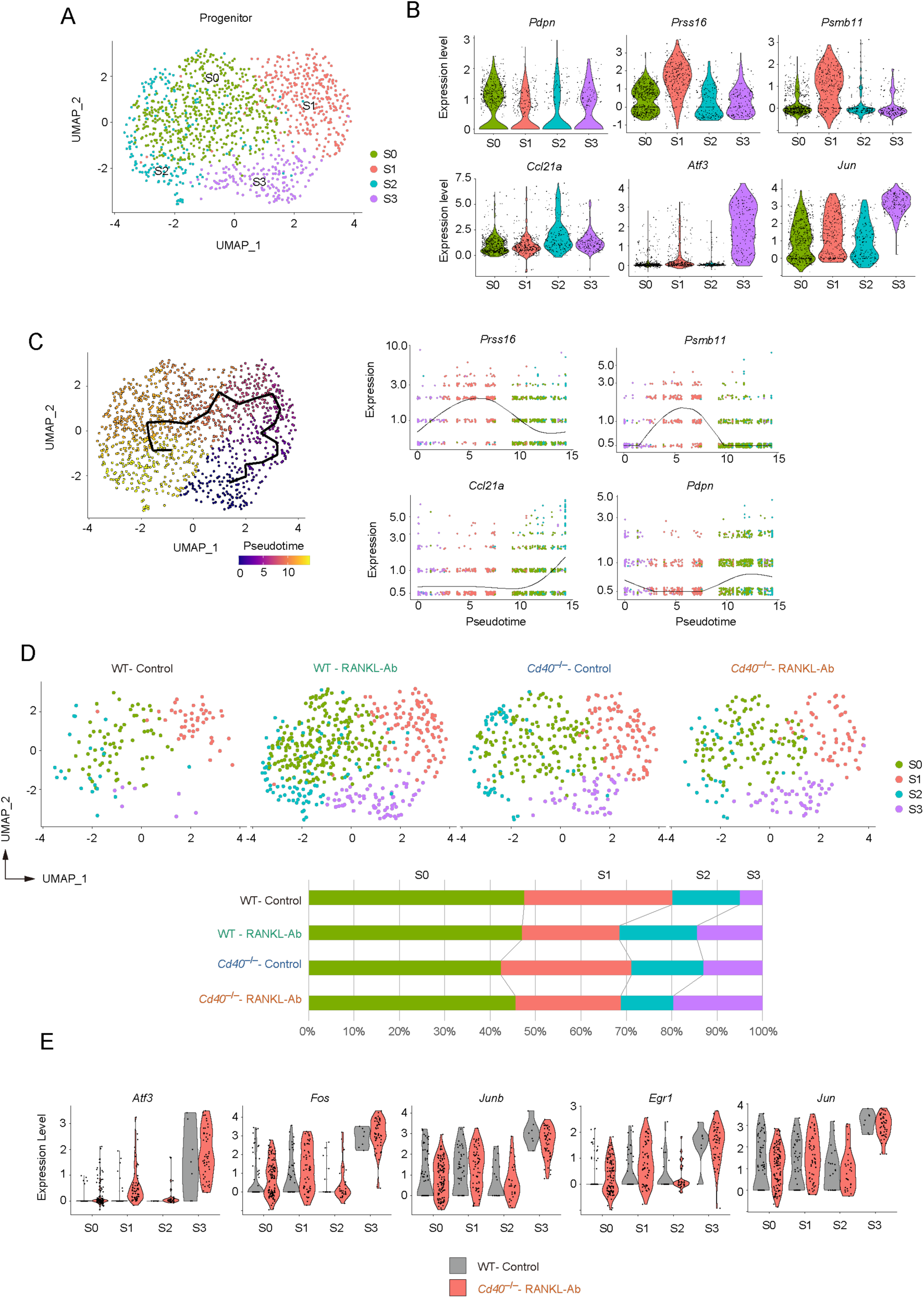
Subclustering analysis of the TEC progenitor subset in scRNA-seq data from wild-type and *Cd40*-deficient mice receiving neutralizing RANKL antibody. (A) UMAP plot showing TEC progenitor clusters in the droplet-based scRNA-seq data. (B) Violin plots depicting expression levels of *Pdpn*, *Prss16*, *Psmb11*, *Ccl21a*, *Atf3*, and *Jun* across each subcluster of TEC progenitors. Each dot represents the expression level in individual cells. (C) The left panel shows a trajectory analysis of subclusters predicted by Monocle 3, visualized by UMAP. Cells are color-coded by pseudotime, transitioning from purple to yellow as pseudotime progresses. The right panel displays the dynamics of gene expression along pseudotime for *Prss16*, *Psmb11*, *Ccl21a*, and *Pdpn*. (D) The integrated UMAP plot from Figure 5A is separated by dataset. The percentages of each cell cluster across the four scRNA-seq datasets are shown in the graph. (E) Violin plots illustrating the expression levels of *Atf3*, *Fos*, *Junb*, *Egr1*, and *Jun* in each subcluster for WT-control and *Cd40*^−/−^-RANKL-Ab. Each dot represents the expression level in an individual cell.

To understand the lineage connections among these clusters, we applied Monocle trajectory analysis to the progenitor cluster. Interestingly, the trajectory analysis suggested an ordering of the clusters in the sequence S3, S1, S0, and S2 (Figure 5C). Assuming that the S3 cluster represents the root node, the pseudotime analysis indicated that the cTEC-biased cluster S1 may differentiate into the mTEC-biased cluster S2 through the non-biased cluster S0.

Comparing the frequencies of subcluster subsets across WT-control, WT-RANKL-Ab, *Cd40*^−/−^- control, and *Cd40*^−/−^-RANKL-Ab mice suggested that the frequency of the S3 cluster increase additively with the elimination of RANKL and CD40L signaling (Figure 5D). In addition, expression of *Atf3*, *Fos*, *Junb*, *Egr1* in other subclusters including cTEC-biased cluster S1 was increased in *Cd40*^−/−^-RANKL-Ab mice (Figure 5E). Thus, the depletion of RANKL and CD40L signaling increases the expression level of AP-1 family genes and the frequency of subsets expressing AP-1 family genes in TEC progenitors.

### Integrative analysis of droplet-based scRNA-seq, well-based scRNA-seq and flow cytometric analyses suggested that the TEC progenitors are present in Ly51^-^UEA-1^-^TEC and cTEC fractions

Our data indicated that TEC progenitor cells, as identified in scRNA-seq analysis, are divided into four clusters depending on gene expression profile. To further validate these subpopulations, we aimed to correlate the scRNA-seq clusters with mTEC and cTEC surface markers in flow cytometric analysis. To this end, we first performed single-cell sorting of UEA-1^+^Ly51^−^ TECs (mTEC-enriched), UEA-1^−^Ly51^+^ TECs (cTEC-enriched), and UEA-1^−^Ly51^−^ TECs (other TECs) from wild-type mouse thymus, followed by RNA-seq of the individual sorted cells (Figure 6A). We then integrated these well-based scRNA-seq data with the droplet-based scRNA-seq data. After assigning each cluster to typical TEC subsets (Supplementary Figure 5), we determined the cell types of the individual cells sorted by flow cytometric analysis (Figure 6B).

**Figure 6.**
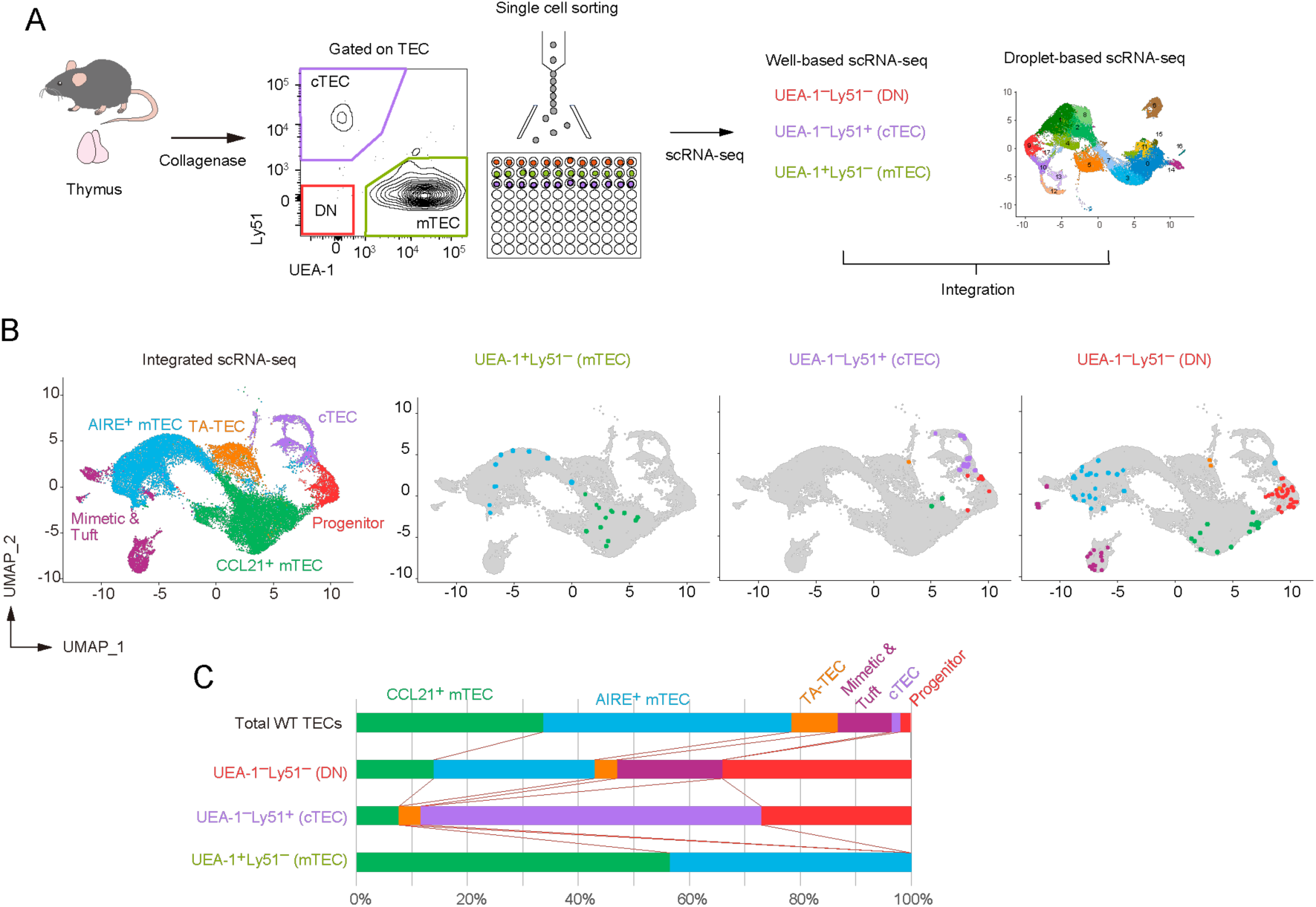
Integration of droplet-based and well-based scRNA-seq data of TECs. (A) Schematic diagram illustrating the integration analysis of well-based scRNA-seq data with droplet-based scRNA-seq data. Single cells were sorted from UEA-1⁺Ly51⁻ TECs (mTECs), UEA- 1⁻Ly51⁺ TECs (cTECs), and UEA-1⁻Ly51⁻ TECs (DN) fractions of wild-type 6-week-old mice, and subjected to well-based scRNA-seq analysis. The well-based scRNA-seq data was integrated with the droplet-based scRNA-seq data shown in Figure 4. (B) UMAP plot and clustering after the integration of droplet-based and well-based scRNA-seq data. Cell clusters were reassigned based on marker gene expression following the integration (Supplementary Figure 4). UMAP plots of individual cells sorted from UEA-1⁺Ly51⁻ TECs (mTECs), UEA-1⁻Ly51⁺ TECs (cTECs), and UEA-1⁻Ly51⁻ TECs (DN) were overlaid on the droplet-based scRNA-seq data (in gray). (C) Percentages of TEC subsets among total single cells in UEA-1⁺Ly51⁻ TECs (mTECs), UEA- 1⁻Ly51⁺ TECs (cTECs), and UEA-1⁻Ly51⁻ TECs (DN).

As expected, our analysis revealed that the UEA-1^+^Ly51^−^ mTEC fraction includes both CCL21^+^ mTECs and AIRE^+^ mTECs (Figure 6B). Notably, the UEA-1^−^Ly51^−^ TEC fraction contains approximately 30% of cells classified as TEC progenitors (Figure 6C), along with some contamination from various mTEC subsets, likely due to the loss of UEA-1 binding ligands during TEC sample preparation using collagenase digestion. Additionally, the UEA-1^−^Ly51^+^ cTEC fraction also contains TEC progenitors alongside mature cTECs. Overall, our data suggest that the TEC progenitors identified in scRNA-seq analysis are negative for UEA-1 binding ligands and are further distinguished based on Ly51 expression levels in flow cytometric analysis.

We next assigned sorted individual progenitor cells to the subpopulations of TEC progenitors identified through droplet-based scRNA-seq analysis. Data analysis revealed that the UEA-1^−^Ly51^−^ TEC fraction contains all types of the TEC progenitor subpopulation (Figure 7A). In contrast, with one exception, progenitor cells sorted from the UEA-1^−^Ly51^+^ TECs predominantly belong to the cluster S1 (Figure 7A), which showed high expressions of cTEC genes (Figure 5A).

**Figure 7.**
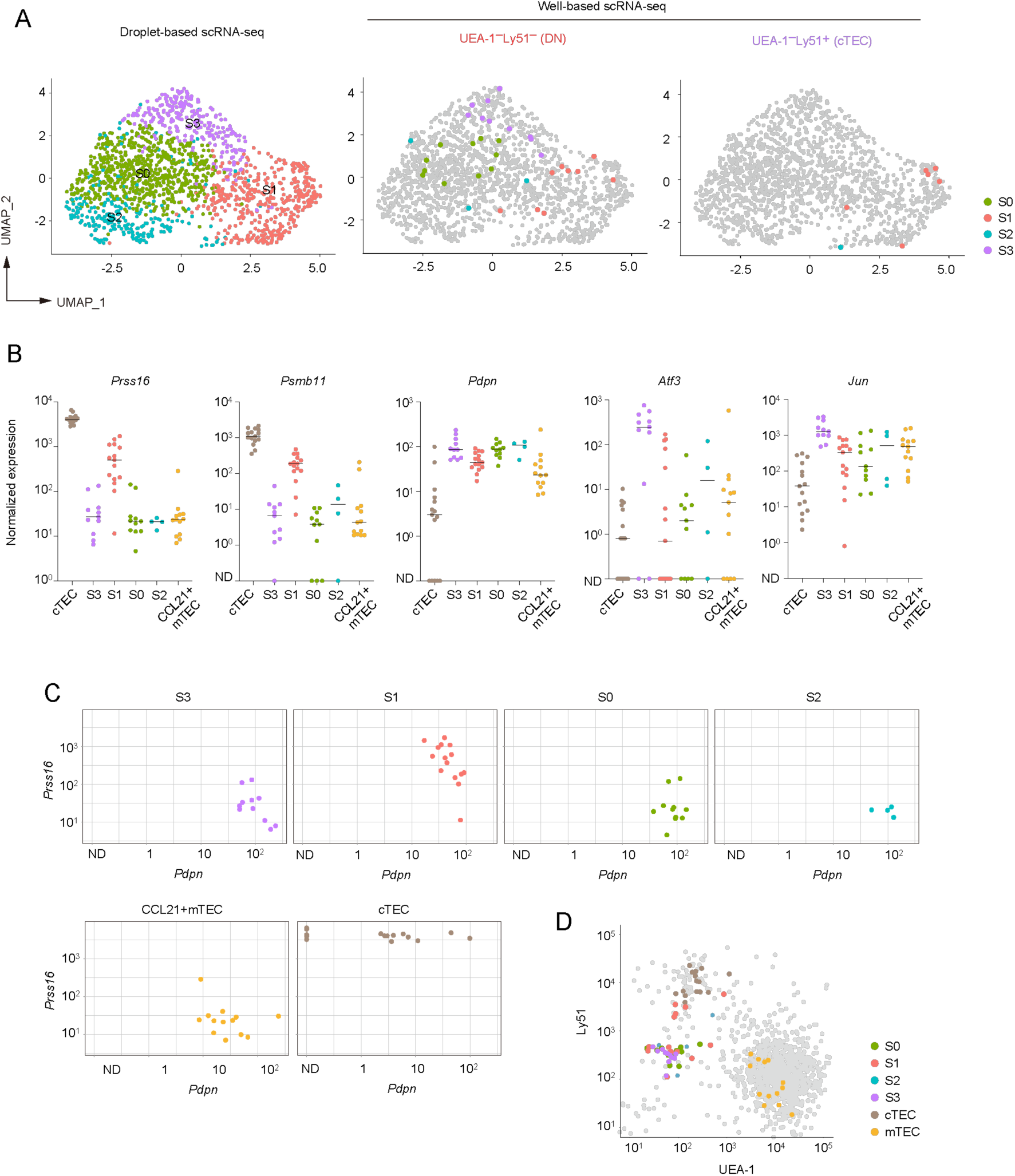
Subclustering analysis of TEC progenitors in integrated scRNA-seq data. (A) UMAP and subclustering of TEC progenitors in the integrated data of the droplet-based scRNA-seq and the well-based scRNA-seq. (B) Dot plots of the normalized expression values of *Prss16*, *Psmb11*, *Pdpn*, *Atf3*, and *Jun* for each cell cluster. Horizontal lines show the median. (C) Scatter plots showing normalized expression values of *Pdpn* and *Prss16* in individual cells of each cluster. One dot represents one cell, color-coded for each cell type. (D) Scatter plot showing the fluorescence intensity of UEA-1 ligand and Ly51 for individual sorted cells assigned as TEC progenitor cells. One dot represents one cell, color-coded for each cluster. Total TECs are indicated as gray dots.

We further investigated the expression level of the cTEC-associated genes and others in the sorted individual cells. In consistent with the droplet-based scRNA-seq data, expression levels of *Prss16* and *Psmb11* were highest in individually sorted cells assigned as the S1 subpopulation (Figure 7B). However, their expression levels were remarkably lower as compared to those in mature cTECs. Expression of *Pdpn* was detected in all subpopulations with almost the same level and may be slightly higher than that in CCL21^+^ mTECs (Figure 7B). Sorted single cells assigned as the cluster S3 exhibited high levels of *Atf3* and *Jun* expressions. In contrast, *Atf3* and *Jun* expression levels were lower in other subpopulations in addition to CCL21^+^ mTECs (Figure 7B). Plot of expression levels of *Pdpn* and *Prss16* further confirmed the presence of the progenitor cells expressing both these two genes in cluster S1 (Figure 7C) in addition to subpopulations expressing only *Pdpn.* These data further confirmed that TEC progenitors are subdivided by expression levels of some AP-1 genes and cTEC-associated genes.

Finally, we confirmed the distribution of these progenitor subpopulations in the flow cytometric profile. As expected, subpopulations S0, S2, S3, and part of S1 were derived from the UEA-1^−^Ly51^+^ fraction and could not be distinguished based on the expression level of these markers (Figure 7D). Interestingly, cells in the S1 cluster within the UEA-1^−^Ly51^+^ TEC fraction exhibited lower surface Ly51 expression compared to cells classified as mature cTECs. This finding suggests that part of the cTEC-biased subpopulation of TEC progenitors is present within the Ly51^lo^UEA-1^−^ fraction in flow cytometric analysis.

## Discussion

DEG analysis of scRNA-seq data suggested that the depletion of RANKL and CD40L signaling influences the gene expression profile not only in RANK-expressing CCL21^+^ mTECs, but also in cTECs and TEC progenitors. Some interferon-stimulated genes were commonly down-regulated in these cell types in *Cd40*^−/−^-RANKL-Ab mice. Type I and III interferons were reportedly expressed in a part of AIRE^+^ mTECs, thereby influencing phenotypes of thymic antigen-presenting cells such as conventional dendritic cells ^35^. Consequently, these interferon signaling also could affect gene expression profiles in cTECs and TEC progenitor cells, suggesting the presence of intercellular communications between mTEC and both cTEC and TEC progenitors, which may affect TEC development ^36^. Our data also suggested an indirect signaling that suppresses the up-regulation of cTEC genes. The mechanism behind this regulation remains to be identified in the future.

We found that the deletion of RANKL and CD40L signaling upregulates AP-1 transcription factor genes selectively in TEC progenitors. Sub-clustering analysis revealed an increase in a TEC progenitor subpopulation that expresses high levels of AP-1 transcription factor genes, as well as upregulation of these genes in certain TEC progenitor subpopulations. This suggests that RANKL and CD40L signaling homeostatically regulate both the subpopulation frequency and gene expression of TEC progenitors via an indirect mechanism. Given the severe defects caused by the depletion of RANKL and CD40L signaling, it is plausible that, in addition to interferon signaling, mature mTEC signaling may regulate AP-1 expression in TEC progenitors through a feedback mechanism. A previous study showed that Fos expression driven by the H2-Kb promoter leads to thymic hyperplasia via TEC expansion ^28^, suggesting that increased AP-1 gene expression could enhance TEC development. Disruption of this indirect signaling may increase AP-1-expressing TEC progenitors potentially aiding in the recovery of CCL21^+^ mTEC precursors.

A previous study suggested that *Pdpn*-expressing TECs are localized at the cortico-medullary junction of the thymus, where they are referred to as junctional TECs (jTECs) ^8^. Additionally, RNA-seq analysis in that study revealed that jTECs express *Ccl21a* ^8^. Consistent with these findings, our scRNA-seq analysis showed that CCL21^+^ mTEC clusters express *Pdpn*. Notably, both our scRNA-seq analysis and previous studies have demonstrated that *Pdpn* is expressed in TEC progenitors ^19^. Sub-clustering analysis further revealed that *Pdpn* is present in all subpopulations of TEC progenitors. Together, these results suggest that *Pdpn* marks not only the CCL21^+^ mTEC precursor pool but also TEC progenitors.

Our study suggested that TEC progenitors were separated into four subpopulations. The trajectory study proposed that TEC progenitors expressing a high level of some AP-1 genes may differentiate into a cTEC-biased subpopulation and subsequently into an mTEC-biased subpopulation. A previous study suggested that adult mTECs differentiate from Psmb11-negative TEC progenitors ^37^. Furthermore, a recent study defined this TEC progenitor as an early TEC progenitor based on this observation ^19^. However, our single-cell study revealed that the expression level of Psmb11 in TEC progenitors was approximately ten times lower than that in cTECs. This suggests that Psmb11^lo^Pdpn^+^ TEC progenitor subpopulation could contribute to maintaining the frequency of adult mTECs. A fate-mapping study using specific marker genes in this subpopulation would be crucial for addressing this issue. Ultimately, our findings illuminate the crucial roles of redundant RANKL and CD40L signaling in maintaining mTEC frequency and TEC progenitor properties in the postnatal thymus, offering promising avenues for developing strategies to address thymic hypofunction associated with aging and various stressors.

## Materials and Methods

### Mice and antibody treatment

Female wild-type C57BL/6 mice, aged 3-4-weeks-old, were purchased from CLEA Japan. *Cd40*-deficient mice were established on a C57BL/6 background. All mice were maintained in standard controlled conditions with a 12-h lighting cycle and access to chow and water ad libitum, housed under specific pathogen-free conditions and handled in accordance with Guidelines of the Institutional Animal Care and Use Committee of RIKEN, Yokohama Branch (2018-075). Rat IgG-Isotype Control antibody (abcam, R&D Systems) or Anti-mouse RANK ligand neutralizing antibody (Anti-RANKL antibody: Oriental enzyme) was injected subcutaneously at 5 mg/kg into C57BL/6J background wild-type mice or *Cd40*-deficient mice.

### Isolation and flow cytometric analysis of TECs from mice

Mice were sacrificed using CO2, and thymi were dissected and placed into cold 1× PBS. Adhering non-thymus tissue was carefully cleared off using sharp tweezers under a fluorescence stereomicroscope. Thymi were minced with a razor blade and pipetted up and down in 1 mL of RPMI 1640 (Wako) to remove lymphocytes. Then, thymic fragments were digested in RPMI 1640 containing Liberase (Roche, 0.05U/mL) and DNase I (Sigma-Aldrich, 0.01% w/v) by incubating three times at 37°C for 12 min each. The supernatant was collected, added to 2 mL of FACS buffer (D-PBS (-) with 2% FBS) containing 1 mM EDTA, and centrifuged at 1500 rpm for 5 min. The supernatant was removed and suspended in FACS buffer. After filtering through a 67-µm nylon monofilament mesh, the resulting cell suspension was incubated with anti-mouse CD16/32 in FACS buffer to block nonspecific binding. For flow cytometric analysis and bulk RNA-seq, cells were stained with primary antibodies (anti-CD45, -TER119, -EpCAM, -Ly51, -L1CAM, -I-A/I-E, -CD104, -Ly-6D, biotinylated UEA-1) in FACS buffer and sequentially incubated with secondary reagent (Streptavidin) in FACS buffer. Dead cells were excluded by staining with SYTOX™ Blue. For droplet-based scRNA-seq, cells were stained with antibodies (anti-CD45, -TER119, -EpCAM) in FACS buffer. For well-based scRNA-seq, cells were stained with primary antibodies (anti-CD45, -TER119, -EpCAM, -Ly51, biotinylated UEA-1) in FACS buffer and depleted of hematopoietic cells and erythrocytes by Magnetic-Activated Cell Sorting (MACS) using APC-MicroBeads (Miltenyi Biotec). Then, cells were stained with secondary reagent (Streptavidin) in FACS buffer. Dead cells were excluded by staining with 7-Aminoactinomycin D. Cells were sorted using a FACS Aria instrument (BD). Data were analyzed using Flowjo 10.

### Bulk RNA-seq analysis

Cells were sorted using a cell sorter (Aria; BD) into 1.5 ml tube with 20 μL of cell lysis solution (2xTCL, 2-Mercaptoethanol). Cell lysis solution or RNase-free water was added to the sorted sample to achieve the final 1x TCL, mixed using a vortex, and the mixture was kept on ice for 5 min. The mixture was centrifuged at 13,000 rpm for 1 min and then stored at -80°C. Cell lysate was dissolved on ice and then purified using total x2.2 volumes of RNAClean XP Beads using Magna Stand. The mixtures were eluted with 40U of RNasin® Plus Ribonuclease Inhibitor in RNase-free water. The supernatant was collected using Magna Stand and denatured at 65°C for 5 min. The mixture was rapidly cooled on ice for 2 min, and then added 10 μL of DNase I solution (PrimeScript Buffer and 2U of DNase I, Amplification Grade in RNase-free water). The mixtures were incubated in a thermal cycler at 30°C for 15 min. Ten μL of first strand cDNA synthesis solution (PrimeScript Buffer, PrimeScript RT Enzyme Mix I, 1 μg of T4 Gene 32 Protein, 6 pmol Oligo(dT)18 Primer and 100 pmol 1st-NSR primer) was added to the DNase I-treated mixture. The mixtures were incubated in a thermal cycler at 25°C for 10 min, 30°C for 10 min, 37°C for 30 min, 50°C for 5 min and 94°C for 5 min. Twenty μL of second strand cDNA synthesis solution (NEBuffer™ 2, 0.625mM dNTP Solution Mix, 500 pmol 2nd-NSR primer and 6.5 U of Klenow Fragment (3’→5’ exo-) in RNase-free water) was added to the first strand cDNA lysate. The mixtures were incubated in a thermal cycler at 16°C for 60 min, 70°C for 10 min. The mixtures were purified with 100 μL of AMPure XP SPRI beads (Beckman Coulter) using Magna Stand, and the concentration were then quantified using Qubit™ dsDNA Quantification Assay Kits. Of the purified dsDNA, 1 ng was used for library preparation, and the rest was stored at -80°C. Thirty μL of tagmentation solution (10 mM Tris-HClpH 8.5, 5 mM MgCl2, 10% N, N-Dimethylformamide and Tn5-linker complex in RNase-free water) and incubated at 55°C for 10 min. Zero-point two percent SDS were added to the mixture and incubated at room temperature for 5 min. Then, the mixtures were purified using Monarch® PCR & DNA Cleanup Kit and eluted into 15 μL of buffer EB. After ligation of adapters using PCR on 25 µl of the purified mixture, sequencing library DNA was purified with x1.2 volumes of AMPure XP SPRI beads and eluted into 15 µL of buffer EB. The sequencing library was sequenced in multiplex on the HiSeqX_Ten platform. FASTQ files were processed using Fastp (Chen et al., 2018) and then quantified for annotated genes using CLC Genomics Workbench (Version 21.0.6, QIAGEN). Differential expression analysis was performed using Proportion-based Statical Analysis on CLC (Version 23.0.4).

### Droplet-based scRNA-seq analysis

For scRNA-seq analysis, cell suspensions of thymi from three mice were prepared and pooled for each individual scRNA-seq experiment. Cellular suspensions were loaded onto a Chromium instrument (10× Genomics) to generate a single cell emulsion. scRNA-seq libraries were prepared using Chromium Next GEM Single Cell 3ʹ GEM, Library & Gel Bead Kit v3.1 and sequenced in multiplex on the HiSeqX Ten platform. FASTQ files were processed using Fastp. Reads were demultiplexed and mapped to the mm10 reference genome using Cell Ranger (v 5.0.1). Processing of data with the Cell Ranger pipeline was performed using the HOKUSAI supercomputer at RIKEN and the NIG supercomputer at ROIS National Institute of Genetics. Expression count matrices were prepared by counting unique molecule identifiers. Downstream single-cell analyses (integration of datasets, correction of dataset-specific batch effects, UMAP dimensional reduction, cell cluster identification, conserved marker identification, and regressing out cell cycle genes) were performed using Seurat v4. Genes that were expressed in more than five cells and cells expressing at least 200 genes were selected for analysis. Cells that contained a percentage of mitochondrial transcripts greater than 13% to 25% were filtered out. Four scRNA-seq datasets were integrated with a combination of Find Integration Anchors and Integrate Data functions. Resolution was set as 0.43 for the FindClusters function. Murine cell cycle genes equivalent to human cell cycle genes listed in Seurat were used for assigning cell cycle scores. Trajectory analysis was performed using Monocle 3.

### Well-based scRNA-seq analysis

Single cells were sorted using a cell sorter (Aria; BD) into 96-well PCR plates with 1 μL of cell lysis solution (1:10 Cell Lysis buffer [Roche], 10 U/μL Rnasin plus Ribonuclease inhibitor [Promega]) in each well, shaken at 1400 rpm for 1 min using a thermo mixer and then stored at -80°C. Cell lysate was dissolved on ice and then denatured at 70°C for 90 sec. To eliminate genomic DNA contamination, 1 μL of genomic DNA digestion solution (PrimeScript Buffer and 0.2 U of DNase I Amplification Grade in RNase-free water) was added to each denatured sample. The mixtures were shaken at 1400 rpm for 1 min using a thermo mixer, and then incubated in a thermal cycler at 30°C for 5 min and held on ice until the next step. One μL of first strand cDNA synthesis solution (PrimeScript Buffer, 8 pmol 1st-NSR primer, 0.6 pmol Oligo(dT)18 Primer, 100 ng of T4 gene 32 protein and PrimeScript RT Enzyme Mix I in RNase-free water) was added to each digested lysate. The mixtures were shaken at 1400 rpm for 1 min using a thermo mixer, and then incubated in a thermal cycler at 25°C for 10 min, 30°C for 10 min, 37°C for 30 min, 50°C for 5 min and 94°C for 5 min. Two μL of second strand synthesis solution (NEBuffer™ 2, 0.625 mM dNTP Solution Mix, 40 pmol 2nd-NSR primer and 0.75U Klenow Fragment (3’→5’ exo-) in RNase-free water) was added to each first strand cDNA lysate. The mixtures were shaken at 1400 rpm for 1 min using a thermo mixer, and then incubated in a thermal cycler at 16°C for 60 min, 70°C for 10 min. The mixtures were purified 15 μL of AMPure XP SPRI beads (Beckman Coulter) diluted two-fold with Pooling buffer (20% PEG8000, 2.5 M NaCl, 10 mM Tris-HClpH 8.0, 1 mM EDTA, 0.01% NP40) using Magna Stand. The mixtures were eluted with 3.75 μL of tagmentation solution (10 mM Tris-HClpH 8.5, 5 mM MgCl2 and 10% N, N-Dimethylformamide in RNase-free water). 1.25 μL of diluted Tn5-linker complex was added to the eluate and the mixtures were incubated at 55°C for 10 min, and then One point two five μL of 0.2% SDS was added and incubated at room temperature for 5 min. After PCR for adaptor ligation, sequencing library DNA was purified using AMPure XP SPRI beads and eluted into 25 μL of buffer EB. Reads were demultiplexed and mapped to the mm10 reference genome with STAR. Cells with less than or more than half the average count of reads detected were excluded from the analysis. Integration of well-based scRNA-seq data with droplet-based scRNA-seq data and UMAP dimension were performed using Seurat. Genes that were expressed in more than five cells and cells expressing at least 200 genes were selected for analysis.

### Statistical analysis

Statistically significant differences between mean values were determined using unpaired t-test or one-way ANOVA followed with multiple comparisons by Tukey’s test in GraphPad Prism (* p≦0.05, ** p≦0.01, *** p≦0.001, **** p≦0.0001). Principle component analysis was performed using edgeR package. P-value correction for differential gene expression analysis in bulk RNA-seq was performed using Baggerley’s test on CLC Genomics Workbench.

## Data availability

The bulk RNA-Seq data have been deposited at NCBI Sequence Read Archive under series accession number SUB14671957. scRNA-seq data were deposited at NCBI GEO under accession number GSE276809

## Acknowledgment

This work was supported by Grants-in-Aid for Scientific Research from JSPS (21K19391 and 23K27399 to T.A., 23K06385 to N.A., and 24K18386 to T.M) and by CREST from the Japan Science and Technology Agency (JPMJCR2011 to T.A.). This work was supported by RIKEN Junior Research Associate Program (MH and HI). The authors declare no competing financial interests.

## Author contributions

**Mio Hayama**, Writing, Data curation, Formal analysis, Investigation, Validation, Writing; **Hiroto Ishii,** Writing – review and editing,Data curation, Formal analysis, Investigation, Validation; **Maki Miyauchi**, Writing – review and editing,Investigation, Validation; **Masaki Yoshida,** Writing – review and editing, Investigation; **Wataru Muramtatu,** Writing – review and editing, Investigation, Validation**; Kano Namiki**, Writing – review and editing, Investigation, Validation; **Naho Hagiwara**, Writing – review and editing, Data curation; **Takahisa Miyao**, Writing – review and editing, Investigation, Validation, Funding acquisition, Supervision; **Nobuko Akiyama,** Supervision, Investigation, Funding acquisition Writing – review and editing; **Taishin Akiyama,** Conceptualization, Funding acquisition, Investigation, Project administration, Supervision, Validation, Writing.

---

**Supplementary Figure 1.**
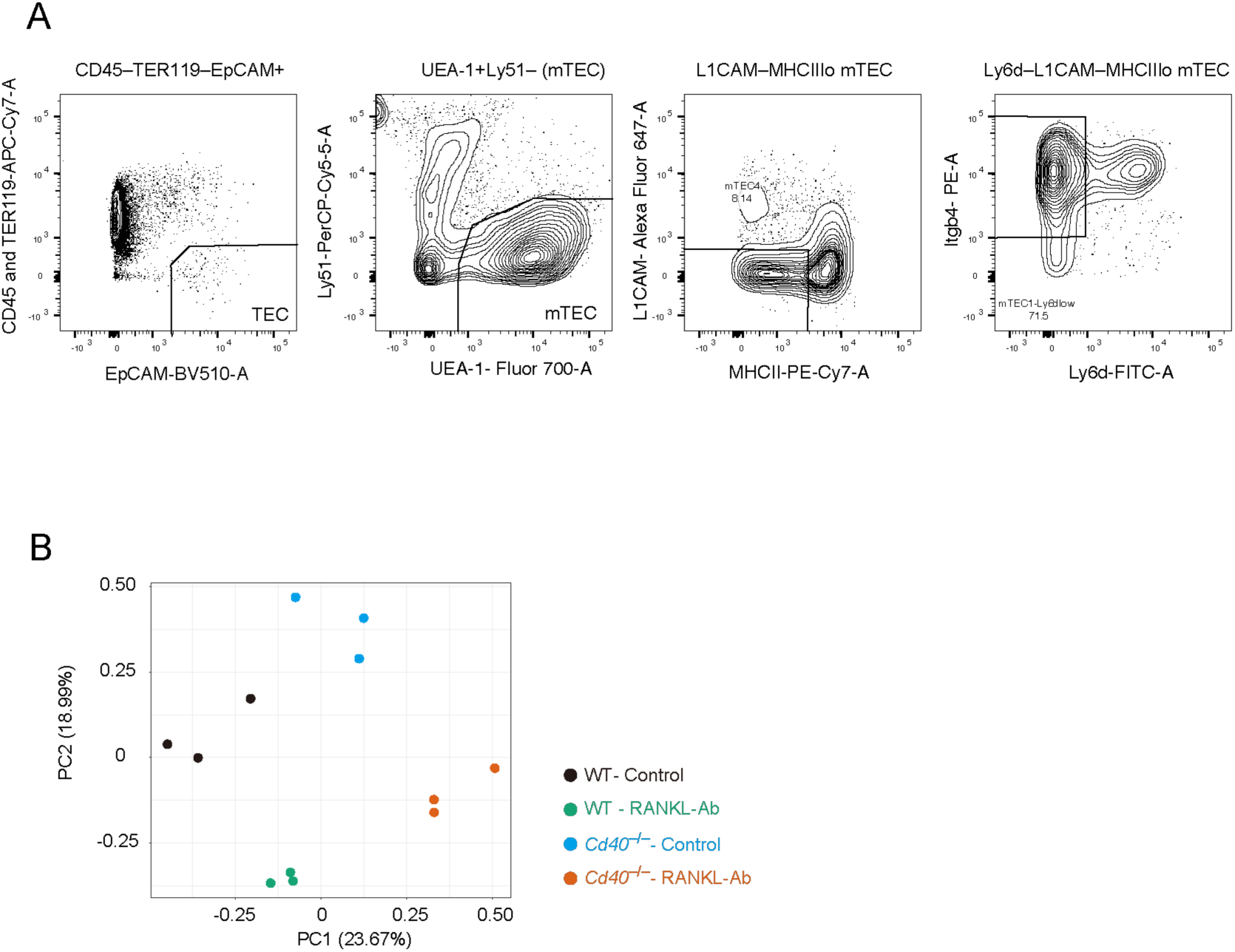
Gating strategy and PCA analysis for bulk RNA-seq analysis of mTEC^lo^ fraction. (A) Gating strategy for sorting the mTEC^lo^ fraction expressing low levels of L1CAM and Ly6d. (B) Principal Component Analysis (PCA) of bulk RNA-seq data from wild-type (WT) treated with control IgG (WT-Control), WT treated with neutralizing RANKL antibody (WT-RANKL-Ab), *Cd40*-deficient (*Cd40*^−/−^) mice treated with control-IgG (*Cd40*^−/−^-Control), and *Cd40*^−/−^ mice treated with RANKL-Ab (*Cd40*^−/−^-RANKL-Ab) at 6-week-old age (n = 3).

**Supplementary Figure 2.**
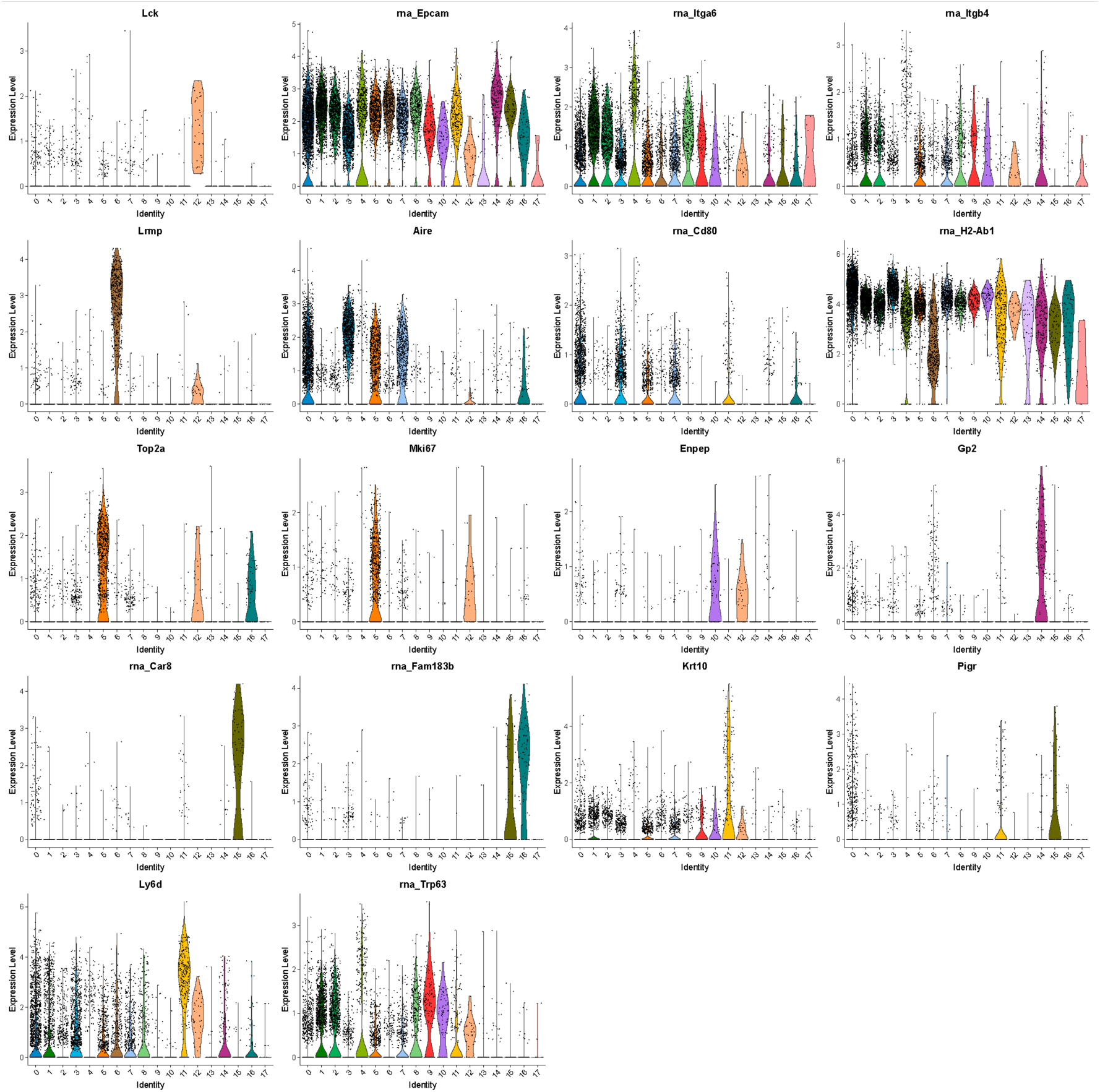
Volcano plot of marker gene expression in cell clusters from scRNA-seq analysis of wild-type mice injected with control IgG. Violin plots showing the expression levels of typical marker genes of TECs in each cluster. The expression levels of these genes in WT treated with control IgG were exhibited. Each dot represents expression levels in individual cells.

**Supplementary Figure 3.**
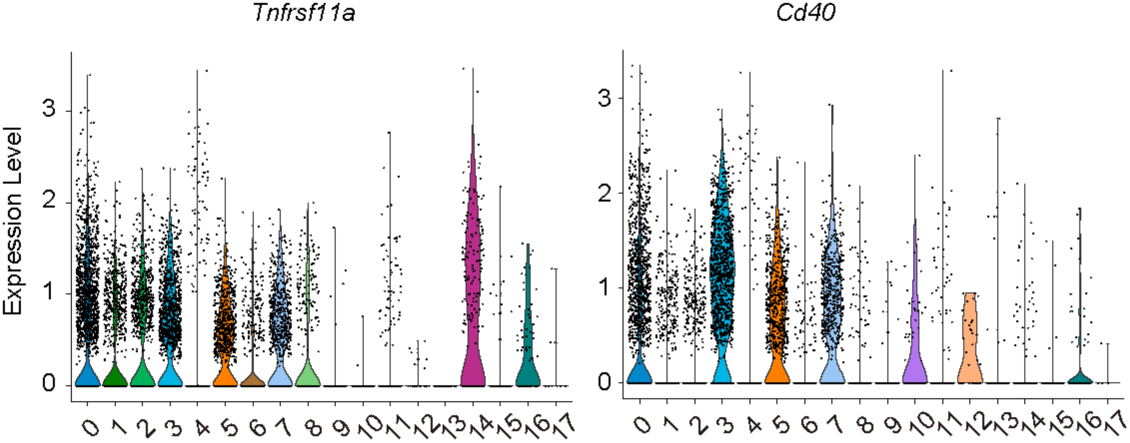
Expression of RANK and CD40 in TEC clusters from scRNA-seq analysis of wild-type mice injected with control IgG. Violin plots showing the expression levels of *Tnfrsf11a (Rank)* and *Cd40* in each cluster. The expression levels of these genes in WT treated with control IgG were exhibited. Each dot represents expression levels in individual cells.

**Supplementary Figure 4.**
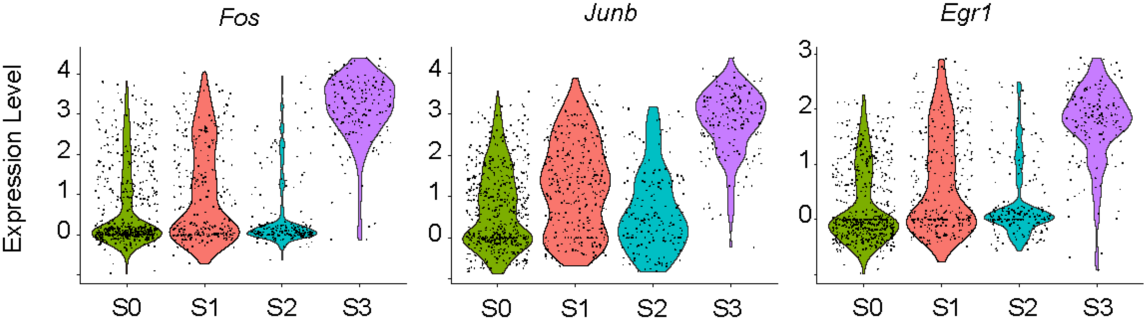
Expression of *Fos*, *JunB*, and *Egr1* in subclusters of TEC progenitors in droplet-based scRNA-seq analysis. Violin plots depicting expression levels of *Fos, Junb* and *Egr1* across each subcluster of TEC progenitors in the droplet-based scRNA-seq data. Each dot represents the expression level in individual cells.

**Supplementary Figure 5.**
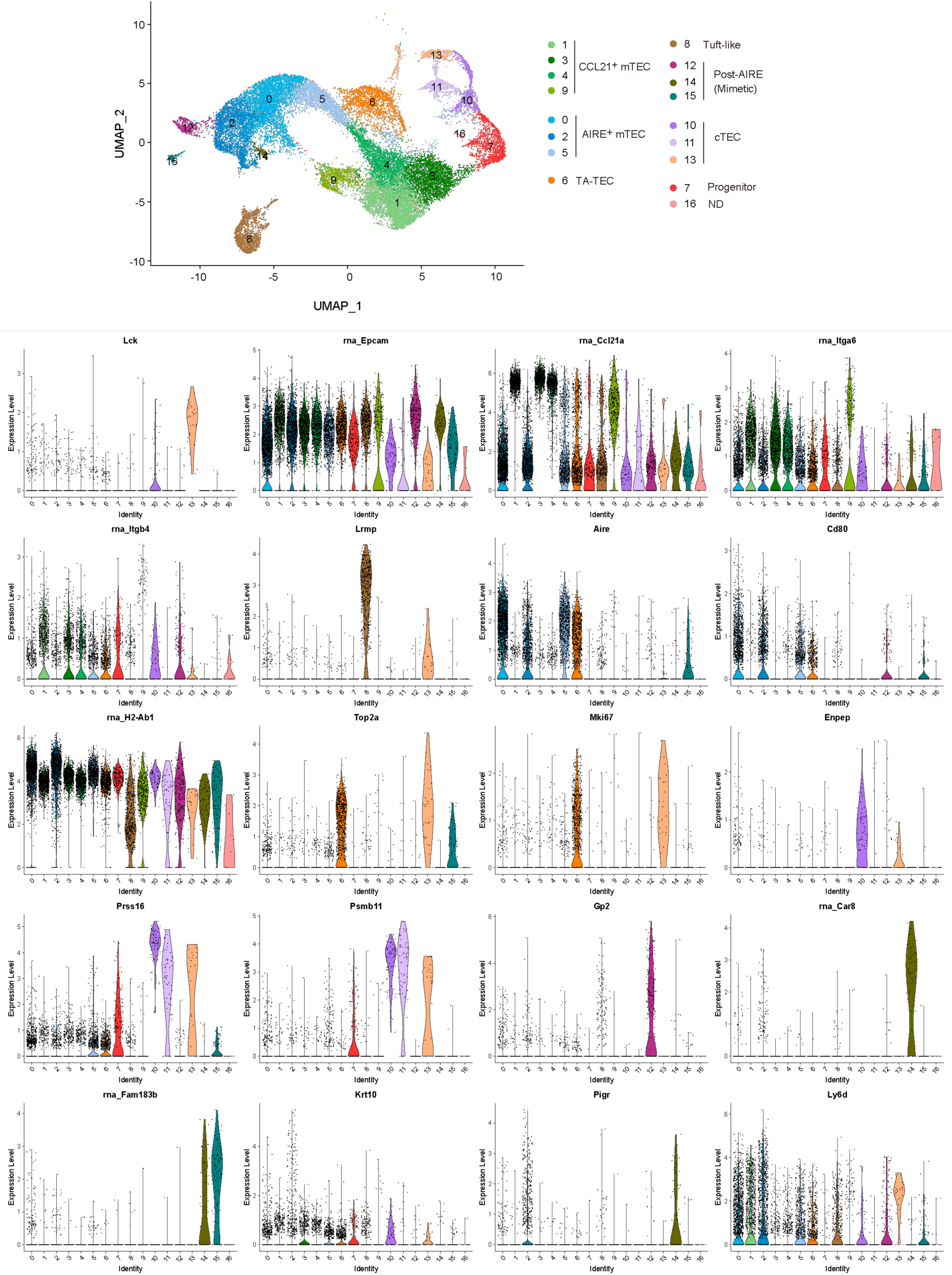
UMAP and clustering of integrated scRNA-seq data combining droplet-based scRNA-seq and well-based scRNA-seq data. The top panel shows UMAP plots after integration of droplet-based scRNA-seq and well-based scRNA-seq using Seurat package. Cell clusters were assigned based on the expression of marker genes. The bottom panels show violin plots showing the expression levels of typical marker genes of TECs in each cluster. The expression levels of these genes in WT treated with control IgG from droplet-based scRNA-seq were exhibited. Each dot represents expression levels in individual cells.

**Supplementary Figure 6.**
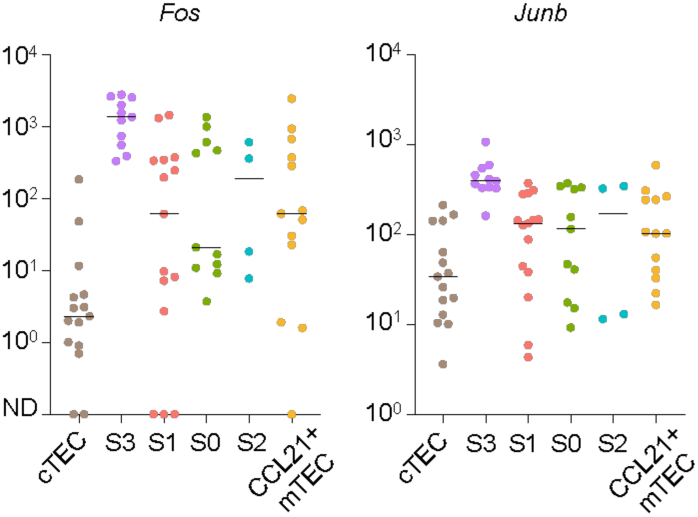
Expression of *Fos* and *Jun* in individual TEC progenitor cells from well-based scRNA-seq analysis. Dot plots of the normalized expression values of *Fos* and *Junb* for each cell cluster. Horizontal lines show the median.

**Supplementary Figure 7.**
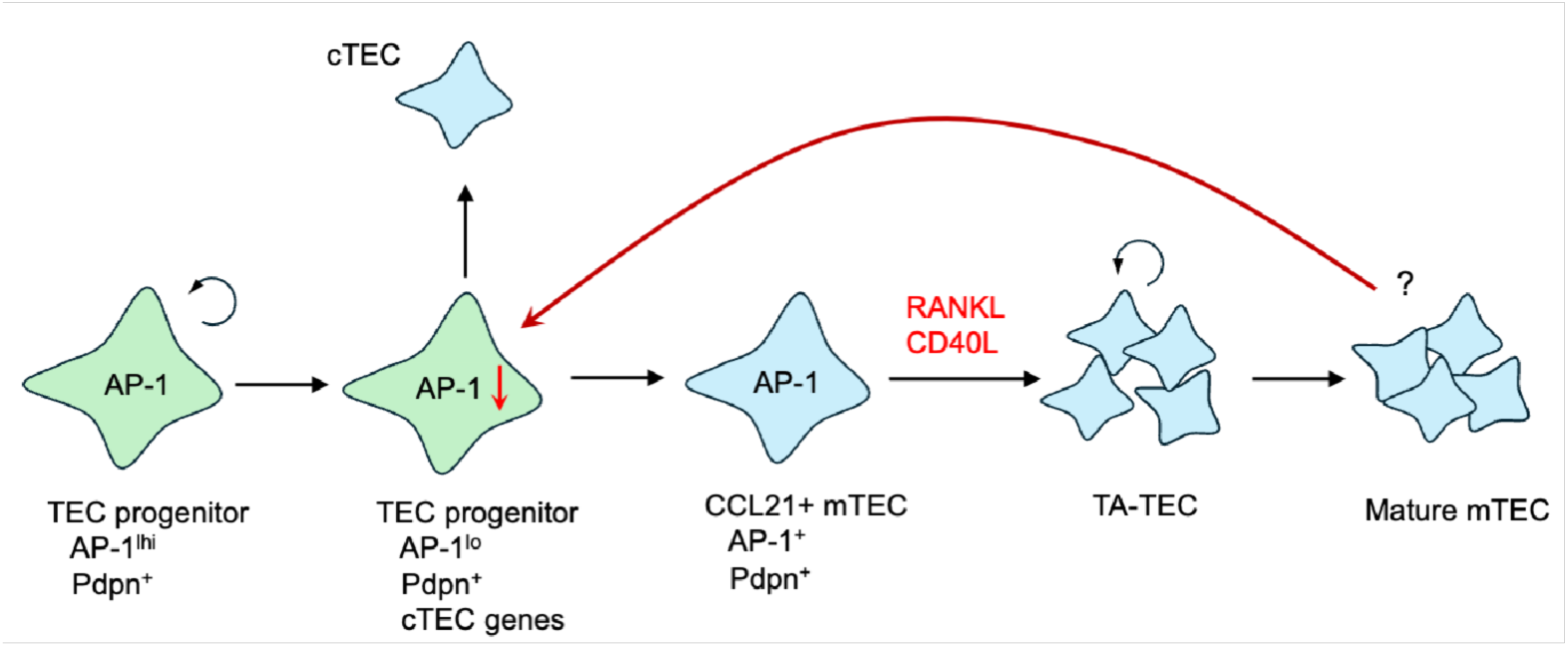
A hypothesis for a feedback regulation of TEC progenitors by indirect RANKL and CD40L signaling. RANKL and CD40L signaling promote the differentiation of CCL21+mTEC into mature mTECs. Mature mTECs control gene expression of TEC progenitors through signaling from mature mTECs. Depletion of RANKL and CD40L signaling results in the elimination of the feedback signaling, thereby up-regulating AP-1 expression in TEC progenitors.

## Notes

### Competing Interest Statement

The authors have declared no competing interest.

